# Transcriptomic profiling of human γδ T cells reveals non-linear immune aging characterized by childhood transitions and relative stability in adulthood

**DOI:** 10.64898/2026.01.15.699622

**Authors:** Tao Yang, Ximena León-Lara, Vicente Almeida, Ziqing Wang, Yusuf E. Abu, Anika Janssen, Charlotte Kleine-Wechelmann, Ahmed Hassan, Zheng Song, Moana Marilyn Dempsey, Likai Tan, Martin Boehne, Constantin von Kaisenberg, Immo Prinz, Reinhold Förster, Sarina Ravens

**Affiliations:** Institute of Immunology, Hannover Medical School, Hannover, Germany; Institute of Systems Immunology, University Medical Center Hamburg-Eppendorf (UKE), Hamburg, Germany; Department of Anaesthesia and Intensive Care and Peter Hung Pain Research Institute, The Chinese University of Hong Kong, Hong Kong SAR, China; Li Ka Shing Institute of Health Sciences, The Chinese University of Hong Kong, Hong Kong SAR, China; Department of Pediatric Cardiology and Intensive Care Medicine, Hannover Medical School, Hannover, Germany; Department of Obstetrics, Gynecology and Reproductive Medicine, Hannover Medical School (MHH), Hannover, Germany; Cluster of Excellence RESIST (EXC 2155), Hannover Medical School, 30625 Hannover, Germany; German Centre for Infection Research, Partner Site Hannover-Braunschweig, Hannover, Germany

## Abstract

γδ T cells are one of the first T cell subsets developing in early ontogeny and show various effector functions in immune homeostasis and response in the young and the old. However, their maturation trajectories from infancy to children, adults and elderly have not been systematically defined. Here, we generated a single-cell transcriptome atlas of 106,711 γδ T cells from 223 individuals spanning infancy to old age. Our analysis reveals that γδ T cell aging is non-linear, characterized by pronounced childhood transitions followed by relative stability throughout adulthood despite marked inter-individual variability. In childhood, changes from developmental and mitochondrial programs toward cytotoxicity and inflammaging were evident. This includes maturation trajectories from GZMK⁺ intermediates to GZMB^+^Perforin^+^ effectors at both RNA and protein levels. Taken together, our study delineates the aging trajectories of human γδ T cells, establishes γδ T cells as a cellular paradigm of non-linear immune aging, and provides a comprehensive resource for investigating γδ T cell biology across the human lifespan.

## Introduction

The traditional view of aging as a gradual, progressive process is increasingly being challenged ^1^. Accumulating evidence supports that aging involves accelerated and non-linear transitions, including multiple biological molecules, age-related brain declines and neurodegenerative diseases such as Parkinson’s disease and Alzheimer’s disease^2–9^. At the cellular level, single-cell transcriptomic profiling has enabled systematic investigation of immune aging but mainly use linear models ^10–12^. Therefore, the extent to which such models capture stage-specific or non-linear aging dynamics of each immune subset remains incompletely understood.

Age is a key determinant of immune system composition and T cell responsiveness ^13^. In infancy, the T cell pool is shaped by the output of new T cells from the thymus and the first exposures to antigens, with a critical need to balance pathogen defense and environmental tolerance^14^. Moreover, for αβ T cells, this phase includes the establishment of immune memory. During adolescence and into adulthood, thymic output and encounters with new antigens decline, while immunological memory increases. This transition reflects a shift from immune development to maintenance, and the establishment of a relatively stable pool of memory and long-lived T cells in adults^13^. In late adulthood and the elderly immune function declines, marked by chronic low-grade inflammation, reduced T cell functionality, and impaired immune regulation, contributing to increased susceptibility to infections, cancer, and other diseases ^15^. While age-related changes in αβ T cells and other immune cell subsets have been extensively studied ^16,17^, the maturation of γδ T cells from infants to elderly remain incompletely understood ^18–21^.

Human γδ T cells represent a distinct subset of T lymphocytes that express a heterodimeric T cell receptor (TCR) composed of a γ-chain and δ-chain. A defining feature of γδ T cells is the association between the usage of specific Vγ and Vδ gene segments, their ontogenetic origin, and their functionality in disease and homeostasis^18^.

In humans, the predominant subset in peripheral blood comprises Vδ2 T cells, which dominate T cell development in the early fetal thymus, where they are pre-committed to various functionalities^22–24^, that include granzymes (e.g. GZMA and GZMK) and cytokine production. Vδ2 T cells express a semi-invariant TCR composed of a Vδ2^+^ δ-chain paired with a Vγ9-JγP^+^ γ-chain, enabling them to recognize small non-peptidic phosphoantigens, metabolites from the isoprenoid biosynthesis pathway, in a butyrophilin-dependent manner ^25^.

Flow cytometric studies have reported a dynamic remodeling of Vδ2 T cell frequencies from birth to adulthood ^26,27^. During the first weeks of life, fetal-derived Vδ2 T cells increase in frequency and absolute number, and acquire the GZMB and Perforin expression ^28,29^. In adulthood, several studies have observed an initial increase of Vδ2 T cells in young adults, and followed a decrease in late adulthood ^26,30–32^.

The second main human γδ T cell subset is Vδ1 T cell subset that pair with a broader range of γ-chains. They first appear in the late embryonic thymus, and dominate γδ T cell generation postnatally ^18,30,33,34^. In the periphery, they are enriched in mucosal and epithelial tissues such as the gut, skin, and lung, and are detectable in peripheral blood, particularly in individuals with chronic viral infections such as cytomegalovirus (CMV)^35^. Flow cytometric studies have reported that the frequency of Vδ1 T cells in peripheral blood remains relatively stable or increases with age^31,32^.

Beyond flow cytometric studies, a recent single-cell transcriptomic study of PBMCs aged 25–85 years analyzed age-associated changes in human γδ T cells ^16^. This study only observed age-related changes of naïve and GZMB⁺ Vδ1 T cell populations and showed no transcriptional age-dependent changes within adulthood. Taken together, these findings raise the possibility that age-associated changes in γδ T cells may not follow simple gradual patterns. In addition, detailed knowledge on the composition in infancy and childhood, and how this compares to adults is not established.

Here, we generated a single-cell transcriptomic atlas of human γδ T cells spanning the entire lifespan by integrating newly generated early-life datasets with selected public scRNA-seq resources. This comprehensive reference map enables systematic investigation of γδ T cell maturation, aging, and functional adaptation across life stages. Importantly, our analysis reveals that γδ T cell aging is not a continuous linear process, but instead is characterized by pronounced changes during early life followed by relative stability across adulthood. Moreover, our dataset is publicly available and provides a unique framework to study γδ T lymphocytes adaption and response capabilities across the human lifespan.

## RESULTS

### Single-cell transcriptome analysis of human γδ T cells from infants to the elderly

The immune response is tightly affected by age. Recently, single-cell transcriptomic technologies have emerged as a powerful approach to dissect immune cell aging at high resolution, yet most studies in peripheral blood have mainly focused on adults, largely due to the limited availability of early-life samples ^16,36,37^. As a result, the aging of γδ T cells was underrepresented due to their low frequency in PBMCs and the limited early-life data.

Here, we aimed to generate a reference map on the transcriptomes of human peripheral blood γδ T cells from newborns to the elderly at steady-state. For this, we performed single-cell transcriptomic analysis of FACS-sorted γδ T cells from infants (n = 5), children (n = 3), and adolescents (n = 6), which together provide critical early-life anchors for the reference map, as well as from young and old adults (n = 14) (**Fig. 1A**, **Supplemental Table 1**). These newly generated datasets were subsequently integrated with public scRNA-seq datasets, yielding a total of 106,711 γδ T cells from 223 individuals aged 1-90 years (**Fig. 1B, Fig. S1A**) ^16,17,38^. Public datasets included either FACS-sorted γδ T cells from children ^38^ or computationally extracted γδ T cells based on TCR gene expression from adult PBMC datasets ^16,39^, as well as large-scale lifespan datasets predominantly composed of adult samples ^17,39^ (**Fig. S1A**).

**Figure 1:**
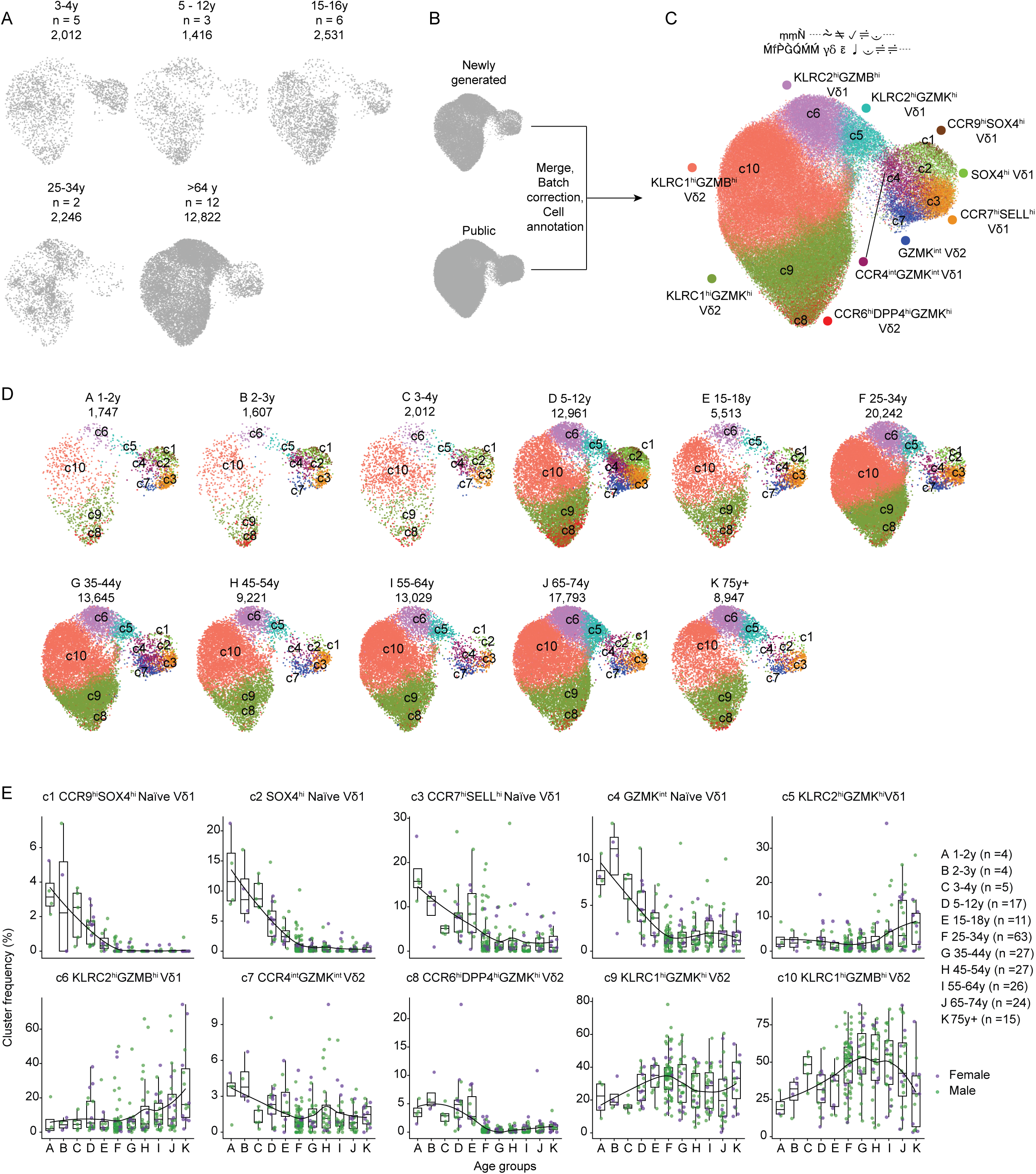
Single-cell transcriptome analysis of human γδ T cells from newborns to elderly. (**A**) Newly generated dataset from this study. Total γδ T cell numbers and donors for each dataset are indicated. (**B**) Workflow for dataset integration, including merging of newly generated dataset, batch correction and cell annotation.. (**C**) UMAP visualization of 106,711 γδ T cells from 225 individuals, colored by cluster identity. (**D**) UMAP plots of γδ T cells from each age group, colored by cluster identity. (**E**) Box plots displaying the frequencies of each cluster across 11 age groups. Each dot represents an individual donor. Boxes indicate median and interquartile range. Male and female donors are indicated. Solid black lines represent LOESS-smoothed trends fitted across age groups.

Following data harmonization across batch and source ^40^, we identified ten γδ T cell clusters annotated based on the expression of TCR usage, transcription factors (TFs), chemokine receptors, and effector molecules (**Fig. 1C, Fig. S1B-C**). Based on *TCF7*, *LEF1*, *GZMA*, *KLRK1* (encoding NKG2D) expression, clusters were categorized into naïve clusters (c1-c4), effector clusters (c5-c6, c8-c10), and a mixed cluster (c7), expressing intermediate levels of *CCR4*, *TCF7*, *GZMA*, and *GZMK* (**Fig. 1C**, **Fig. S1B**). Naïve clusters, which were predominantly Vδ1 T cells and a few Vδ3 T cells, included the following: c1 (*CCR9*^hi^*SOX4*^hi^), c2 (*SOX4*^hi^), c3 (*CCR7*^hi^*SELL*^hi^), and c4 (*GZMK*^int^) clusters. The effector clusters were c5 (*KLRC2*^hi^*GZMK*^hi^ Vδ1-dominant), c6 (*KLRC2*^hi^GZMB^hi^ Vδ1-dominant), c8 (*CCR6*^hi^*DPP4*^hi^*GZMK*^hi^ Vδ2), c9 (*KLRC1*^hi^*GZMK*^hi^ Vδ2), and c10 (*KLRC1*^hi^*GZMB*^hi^ Vδ2).

Next, to examine age-associated compositional trends, all samples were divided into 11 age groups (A – K) (**Fig. 1D-E**). The naïve clusters (c1-c4) and the intermediate cluster c7 decline sharply until puberty (age group E, 15 – 18y), while type-1 immunity effectors (c5-c6, c9-c10) accumulated. The *CCR4*^int^ *GZMK*^int^ cluster c7 had the highest abundance in neonates and infants (age group A – B, 1 – 3 years old), and stayed stable thereafter. The *CCR6*^hi^*GZMK*^hi^ Vδ2 cluster (c8) peaked in childhood and puberty (age group A – E). Thereafter was a rapid decline of the cluster c8 with a stable persistence into elderly individuals. Importantly, above described compositional changes largely occurred before adulthood. From age group F (25y-34y) onward, most clusters stabilized in frequency. As an exception, there was a continued expansion of effector Vδ1 T cells (c5 and c6) with the highest abundance in elderly individuals (age group K, 75y+). *Vice versa*, there was a late-life decline in effector Vδ2 T cells (c9 and c10). Notably, all age-related changes and distribution of clusters were sex-independent (**Fig. S1C**).

In summary, integrating several independent datasets of newly generated and public available single-cell transcriptomes from peripheral blood γδ T cells, we have generated a detailed reference map on the transcriptional programs of peripheral Vδ1 and Vδ2 T cells from infants to elderly. This provides a highly valuable resource to examine maturation trajectories of γδ T cells and to study their transcriptomes in disease and with respect to age.

### Vδ1 T cell maturation bifurcates from a CCR9⁺SOX4⁺ effector precursor subset into migratory and cytotoxic lineages

Next, age-related maturation trajectories were investigated. Building on **Fig. 1E**, the CCR9^hi^SOX4^hi^ Vδ1 cluster (c1) peaked in early infancy and became rare in adulthood. Concordantly, it has been reported the presence of CCR9⁺ γδ T cells in cord blood ^41^ and in the thymus ^34^. Transcriptionally, c1 cells showed exclusive *CCR9* expression, along with high levels of *SOX4*, *TCF7*, *LEF1*, *TCF12*, *ID3*, and *LCK*, all associated with recent thymic emigrants (**Fig. 2A**). Additionally, c1 cells expressed *LAT*, *LTB*, *TRAF3IP3*, *RHOH*, and *CD27*, genes involved in thymocyte survival and lineage commitment. Next, these transcriptional profiles were compared to all of the Vδ1 T cell clusters (c2 – c6). Cluster c2 lacked *CCR9* expression but maintained high expression of above described TFs of Cluster c1. Cluster c3 showed downregulation of all these TFs and upregulation of *CCR7* and *SELL* (encoding CD62L). Notably, c3 also expressed high levels of *FOXP1*, a TF that regulates naïve T cell homing and survival via CCR7/CD62L expression ^42^. Cluster c4 cells express *GZMK*, *GZMA*, *KLRC2, KLRC3*, and *GNLY*, which may indicate the transition toward an effector state (**Fig. 2A**).

**Figure 2:**
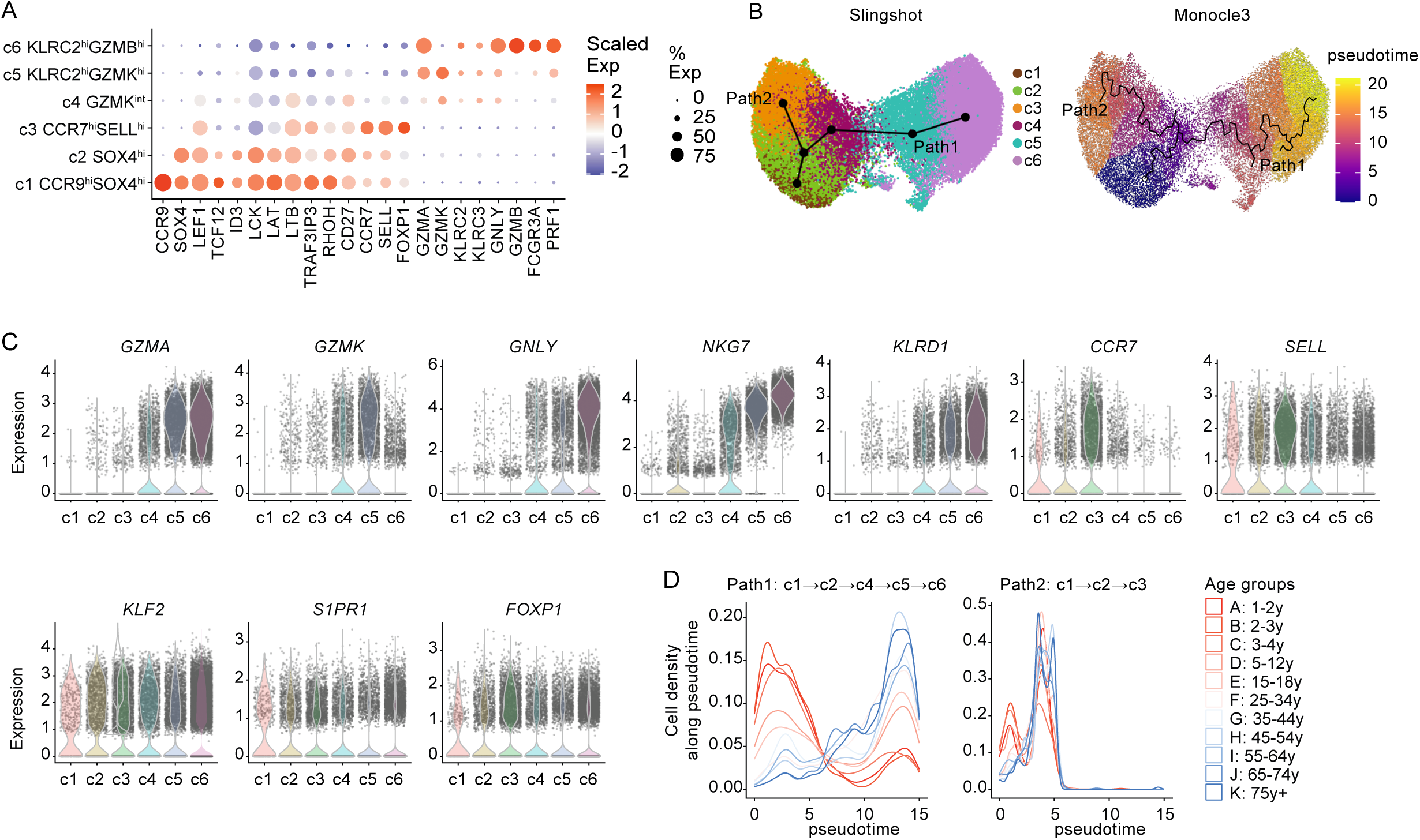
Developmental trajectory analysis of Vδ1 T cells. (**A**) Dot plot showing scaled expression (Scaled Exp) and proportion (% Exp) of cells expressing selected marker genes across clusters c1–c6. (**B**) Pseudotime trajectory inference of Vδ1 T cells using Slingshot (left) and Monocle3 (right). Cells are colored by cluster identity (left) and pseudotime (right), respectively. (**C**) Violin plots showing expression of selected genes across clusters c1–c6. (**D**) Distribution of cells along pseudotime in Path 1 (c1 → c2 → c3) and Path 2 (c1 → c2 → c4 → c5 → c6). Density plots are stratified by age group (A–L); each line represents a group-specific density estimate.

Next, pseudotime analyses using Monocle3 ^43^ and Slingshot ^44^ consistently revealed two divergent differentiation paths originating from the CCR9^hi^SOX4^hi^ naïve Vδ1 cluster (c1): Path-1 (c1 → c2 → c4 → c5 → c6) displayed a stepwise acquisition of cytotoxic programs (e.g., *GZMA*, *GZMK*, *GNLY*, *NKG7*, and *KLRD1*) (**Fig. 2B-C**); Path-2 (c1→ c2 → c3) retained a naïve transcriptional profile (e.g. *TCF7*, *LEF1*, *CD27*) while progressively upregulating tissue-homing and survival-related genes (*CCR7*, *SELL*, *KLF2*, and *FOXP1*) (**Fig. 2A-C**). Along both paths, pseudotime distributions revealed a clear age shift, with cells from younger individuals enriched at early stages and those from older individuals predominating at later stages (**Fig. 2D**).

Together, these findings support a bifurcated peripheral maturation model. The CCR9⁺SOX4⁺ Vδ1 T cells represent a transient population in early life that matures into at least two different fates, one with differentiation into *GZMB*^+^ effector cells in peripheral blood (via c4-c6) and one with a *CCR7*⁺*SELL*⁺ tissue-migratory fate (via c3). Of note, the cytotoxic differentiation trajectory of Vδ1 T cells (Path-1: c1→c2→c4→c5→c6) indicated that *GZMK*⁺ cells in c5 likely represent a transitional pre-effector stage that precedes the fully cytotoxic *GZMB*⁺ population in c6.

### GZMK^+^ γδ T cells represent a pre-effector intermediate population and can differentiate into GZMB^+^Perforin^+^ cytotoxic effectors

GZMB is the most potent inducer of caspase-dependent apoptosis and expressed by terminally differentiated cytotoxic T cells. On the other hand, GZMA and GZMK rather trigger caspase-independent death and promote inflammation^45^. Flow cytometric analysis of PBMCs showed that γδ T cells were either GZMB^+^ or GZMK^+^ (**Fig. 3A**). Notably, GZMK^+^ γδ T cells displayed significantly higher expression of CD27, CD28, CD69, and CD127, and reduced levels of cytotoxic proteins, including CD16 and Perforin (**Fig. 3B**). Hence, we compared the transcriptional profiles of *GZMK*^hi^ clusters (c5, c8, c9) and *GZMB*^hi^ clusters (c6, c10) (**Fig. 3C, Fig. S2A–B**). GZMK^hi^ clusters (c5, c8, and c9) showed elevated expression of *TCF7* and *IL7R* (CD127), and other genes related to pre-effector and memory-like states, including *CD27*, *CD28* (especially enriched in Vδ2^+^ clusters), *LTB*, AP-1 factors (*FOS*, *JUNB*, *JUN*) and *NFKBIA* (**Fig. 3C, Fig. S2A–B**). Conversely, *GZMB*^hi^ clusters (c6 and c10) were enriched for genes linked to a high cytotoxicity such as *PRF1* (Perforin), *GNLY* (Granulysin), *FCGR3A* (CD16), and *KLRD1* (CD94), tissue trafficking (*ITGB1*, *ITGB2*, *ITGAL*), antiviral activity (*IFITM1*, *IFITM2*), and innate activation (*S100A4*, *S100A6*).

**Figure 3:**
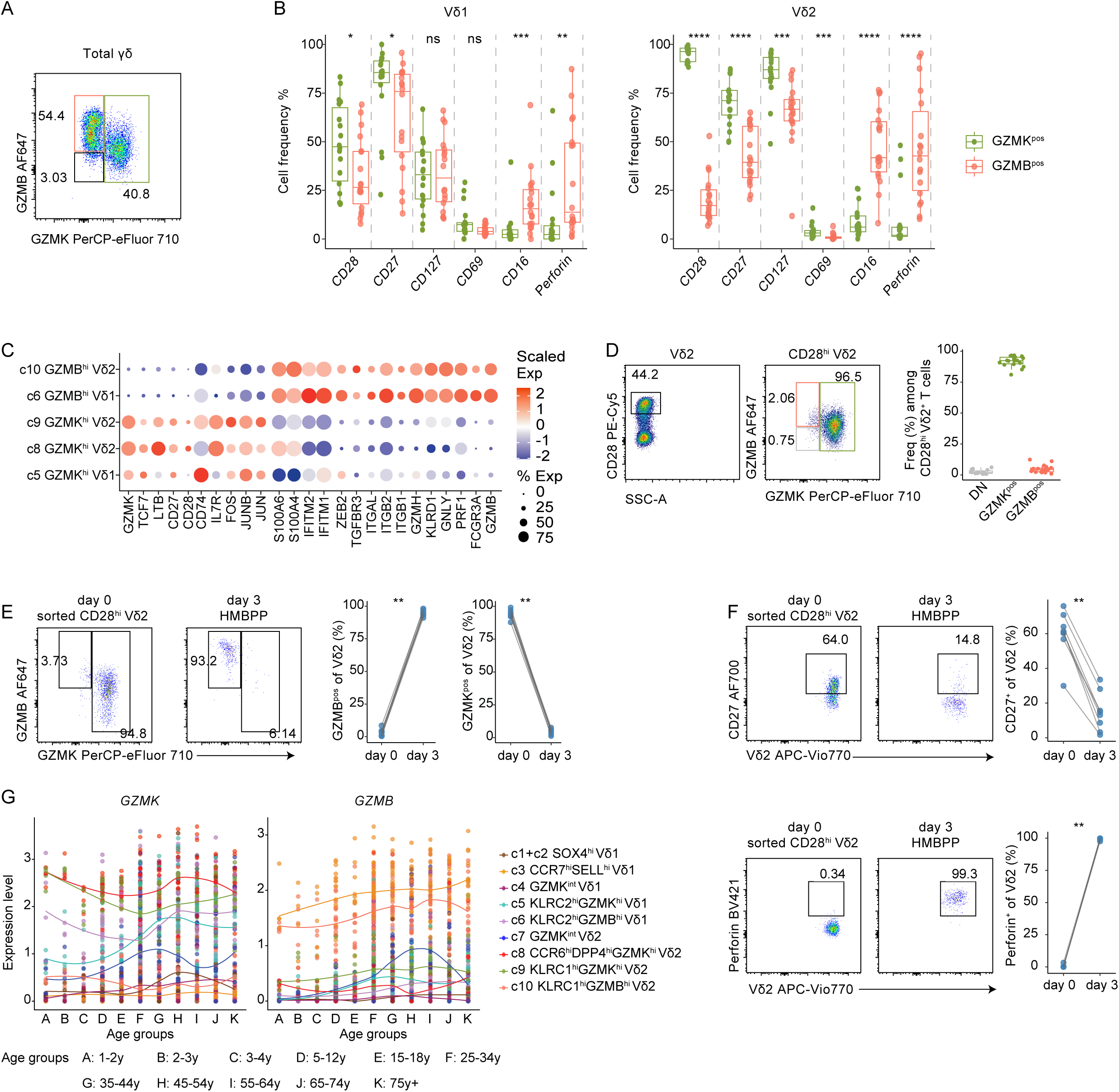
Characterization of GZMK⁺ and GZMB⁺ γδ T cell subsets. (**A**) Representative flow cytometry plot showing GZMK and GZMB expression in total γδ T cells from adult PBMCs. (**B**) Frequencies of CD28, CD27, CD127, CD69, CD16, and Perforin expression in GZMK⁺ and GZMB⁺ Vδ1 (left) and Vδ2 (right) T cells from adult PBMCs (n = 18). Each dot represents one donor. Boxes indicate median and interquartile range. (**C**) Dot plot showing scaled expression (Scaled Exp) and proportion (% Exp) of cells expressing selected marker genes across GZMK⁺ and GZMB⁺ clusters within Vδ1 and Vδ2 T cell subsets. (**D**) Left: Representative flow cytometry plot showing GZMK and GZMB expression within CD28⁺ Vδ2 T cells from adult PBMCs. Right: quantification of the proportion of GZMK⁺ and GZMB⁺ cells within CD28⁺ Vδ2 T cells (n = 18). Each dot represents one donor. Boxes indicate median and interquartile range. (**E**) Induction of GZMB and loss of GZMK in sorted CD28⁺ Vδ2 T cells after 3-day stimulation with HMBPP. Left: representative flow cytometry plots at day 0 and day 3. Right: paired quantification of GZMB⁺ and GZMK⁺ frequencies (n = 10). Lines connect measurements from the same donor. (**F**) CD27 downregulation (top) and Perforin upregulation (bottom) in CD28⁺ Vδ2 T cells following 3-day HMBPP stimulation. Left: representative flow cytometry plots at day 0 and day 3. Right: paired quantification of CD27⁺ and Perforin⁺ cells among Vδ2 T cells (n = 10). (**G**) Expression of GZMK (left) and GZMB (right) across age groups (A–L) and transcriptionally defined γδ T cell clusters, with data points and lines colored by cluster identity. Each dot represents the average expression of the indicated gene for one donor within a given cluster. Solid lines represent LOESS-smoothed trends fitted across age. Statistical comparisons in (**B**) and (**E-G**) were assessed by unpaired two-sided Wilcoxon rank-sum test and paired two-sided Wilcoxon signed-rank test, respectively. Significance is denoted as ns (not significant), *p < 0.05, **p < 0.01, ***p < 0.001, ****p < 0.0001.

CD28 appears to be a reliable surface marker to isolate GZMK^+^ Vδ2 T cells as the majority of those are GZMK⁺ (**Fig. 3D**). Indeed, FACS-sorted adult blood CD28⁺ Vδ2 T cells were predominantly GZMK^+^GZMB^-^ prior to *in vitro* activation (**Fig. 3E**). After three days of *in vitro* (E)-4-hydroxy-3-methyl-but-2-enyl pyrophosphate (HMBPP) stimulation, a decline in GZMK⁺ Vδ2 T cells and a concurrent increase in GZMB⁺ Vδ2 T cells was evident (**Fig. 3E**). This was accompanied by elevated expression of Perforin, and reduced CD27 expression, indicating a maturation of CD28^hi^ GZMK^+^ to GZMB^+^ Perforin^+^ Vδ2 T cells (**Fig. 3F**). Notably, CD28 expression did not reliably distinguish GZMK⁺ from GZMB⁺ populations in Vδ1 T cells, CD8⁺ T cells, or NK cells by flow cytometry (**Fig. S2C**). With regard to age-related transcriptional changes, we show that *GZMK*^hi^ Vδ1 and Vδ2 T cells (c5, c8, and c9) emerged earlier in life than *GZMB*^hi^ Vδ1 and Vδ2 T cells (c6 and c10) (**Fig. 3G**). This age-ordered distribution of *GZMK* and *GZMB* expression in both Vδ1 and Vδ2 T cell lineages suggests an age-associated maturation trajectory from *GZMK*^hi^ to *GZMB*^hi^ γδ T cell effectors in periphery. Notably, reanalysis of a public fetal and pediatric thymus dataset identified *GZMK*^hi^ but not *GZMB*^hi^ γδ T cells at these early developmental stages (**Fig. S2D**)^24^. Flow cytometry further showed: GZMK^+^ γδ T cells were readily detected in cord blood from term neonates, whereas GZMB^+^ γδ T cells were nearly absent (**Fig. S2E**).

Collectively, these results provide transcriptional and phenotypic evidence that GZMK^hi^ γδ T cells represent an intermediate pre-effector population that can differentiate into GZMB^hi^ cytotoxic effectors. This maturation trajectory appears conserved across all γδ T cell subsets.

### Age-associated transcriptional changes in human γδ T cells are characterized by reduced developmental and mitochondrial programs and enhanced effector-related signatures

Next, we systematically investigated gene expression dynamics of γδ T cells across all age groups. To this end, we performed differential gene expression (DEG) analysis between young (age groups A-C) and old (age groups I-K) individuals, identifying 179 upregulated and 335 downregulated genes (**Fig. 4A**).

**Figure 4:**
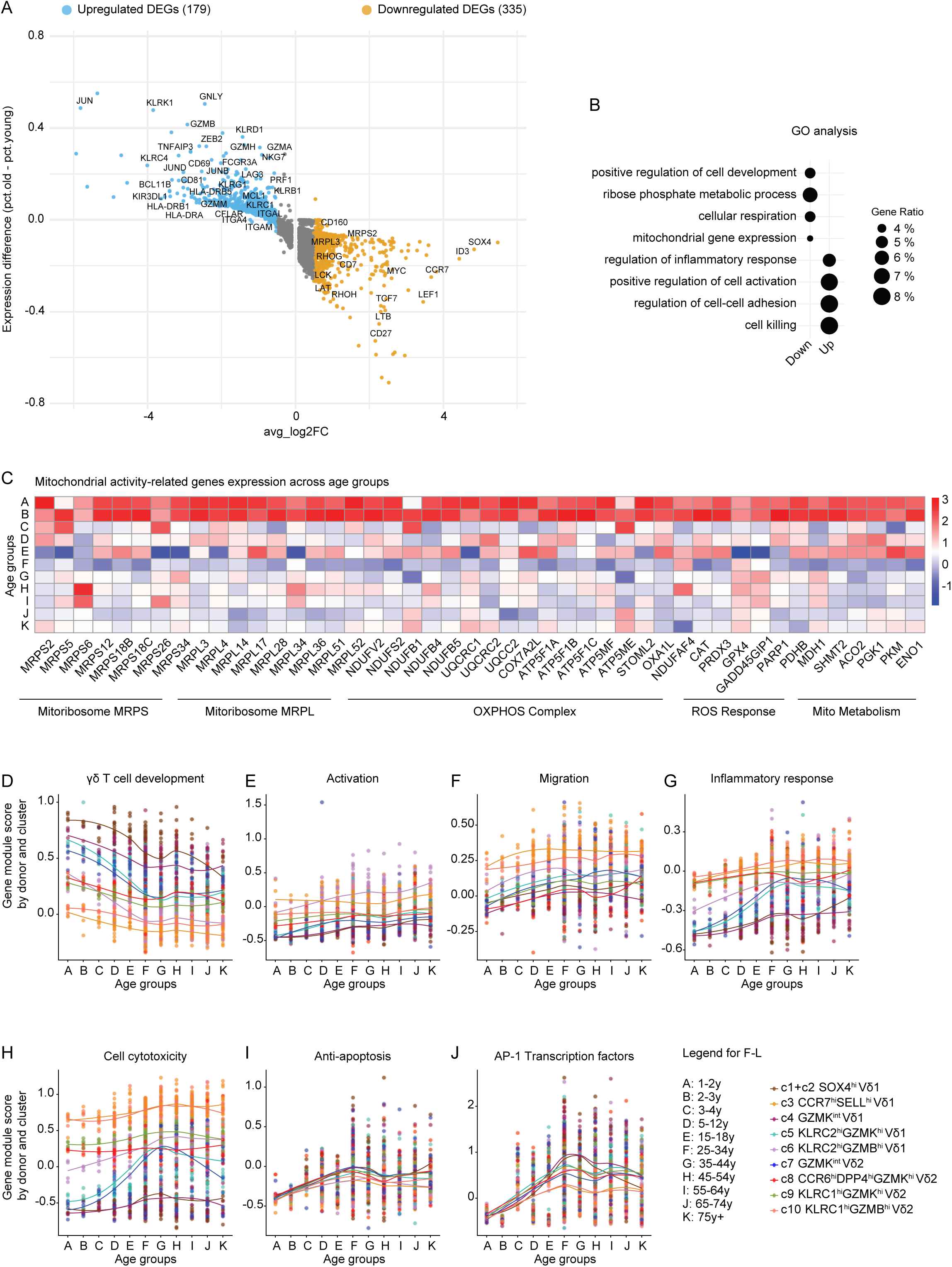
Age-associated transcriptional remodeling and mitochondrial decline in γδ T cells. (**A**) Volcano plot showing average log₂ fold change (x-axis) and expression difference (y-axis) of DEGs in aged versus young γδ T cells. Selected genes are labeled. (**B**) Gene Ontology (GO) enrichment analysis of upregulated and downregulated DEGs. Dot size indicates gene ratio. (**C**) Heatmap showing average expression of mitochondrial activity-related genes across age groups. (**D-J**) Module scores of gene sets related to γδ T cell development (D), activation (**E**), migration (**F**), inflammatory response (**G**), cell cytotoxicity (**H**), anti-apoptosis (**I**), and AP-1 transcription factors (**J**), plotted across age groups and clusters, with data points and lines colored by cluster identity. Each dot represents the average expression of the indicated gene score for one donor within a given cluster. Solid lines represent LOESS-smoothed trends fitted across age. Gene lists used for module scores are shown in Fig. S4.

Gene ontology (GO) analysis revealed that downregulated genes were enriched for mitochondrial functions, including mitochondrial ribosomal small and large subunits (*MRPS* and *MRPL*), TCA cycle enzymes, oxidative phosphorylation (OXPHOS) complexes, and reactive oxygen species (ROS) response genes (**Fig. 4B-C**).

Moreover, age-related declines in genes linked to γδ T cell development were also observed (*LEF1*, *CD27*, *IL7R*, *SOX4*, *TCF7*, *LTB*, *ID3*, *CCR7*, *RHOH*, *LCK*, *RELB*, *RHOG*, *CD7*, *MYC*, *CCR9*, *LAT*) (**Fig. 4A-B**). A composite γδ T cell development score incorporating these genes declined progressively with age (**Fig. 4D**). Conversely, upregulated genes were enriched for effector-related gene expression programs, including activation (*HLA-DRB5*, *CD160*, *CD81*, *ZEB2*, *CD69*), migration (*ITGA4*, *ITGB2*, *CX3CR1*, *CXCR4*, *PREX1*, *TMSB10*), inflammatory response (*NFKBIZ*, *ZBP1*, *TNFRSF1B*, *ALOX5AP*), and cytotoxicity (*PRF1*, *GZMB*, *GNLY*, *KLRK1*) (**Fig. 4A**, **Fig. 4E–H, Fig. S3A**). Anti-apoptotic genes (e.g. *BCL2*, *MCL1*, *CFLAR*) were also elevated, suggesting enhanced survival capacity with age. In addition, AP-1 transcription factors (*JUN*, *JUNB*, *JUND*, *FOS*, *FOSB*, *FOSL2*) were elevated (**Fig. 4I–J**) suggesting a shift toward activation-primed states with age.

To further assess effector potential and immune adaptation of γδ T cells across age, we examined gene signatures, including cytokine receptors, chemokine ligands and receptors, TCR signaling components, toll-like receptors, and proliferation (**Fig. S3B**). In parallel, we evaluated canonical features of immune aging^15^, including T cell exhaustion, senescence-associated secretory phenotype (SASP), DNA repair and genome stability, autophagy, and epigenetic regulation (**Fig. S3C**). These pathways displayed heterogeneous changes: some genes were expressed at very low levels (e.g. *IL1A*, *IL4*), others showed only minor variation with age (e.g. *TGFBR2*, *IFNAR1*), and a small subset was either upregulated or downregulated. For example, reduced *IL23R* and *CXCR6* expression, was consistent with the abundance of type-3 like γδ T cells in adults^41^. Notably, *LAG3* was selectively increased in older individuals, potentially indicating heightened activation with a potentially impaired function ^46^.

Taken together, these findings reveal age-associated transcriptional changes of human γδ T cells across the lifespan. In infancy and childhood, γδ T cells undergo pronounced transcriptional changes, characterized by a reduction in developmental and mitochondrial gene signatures and a concomitant increase in genes related to activation and cytotoxicity, while changes are relatively modest during adulthood. This raised a need to identify a representative way to capture γδ T cell aging across the lifespan.

### γδ T cell RNAge assesses age-related transcriptional programs and captures inter-individual heterogeneity

To integrate age-associated transcriptional changes of γδ T cells, we identified a γδ T cell RNA age score (RNAge) based on gene programs identified in Figure 4 (**Fig. 5A**). γδ T cell RNAge showed a pronounced rise in childhood and relative stabilization thereafter in adulthood.

**Figure 5:**
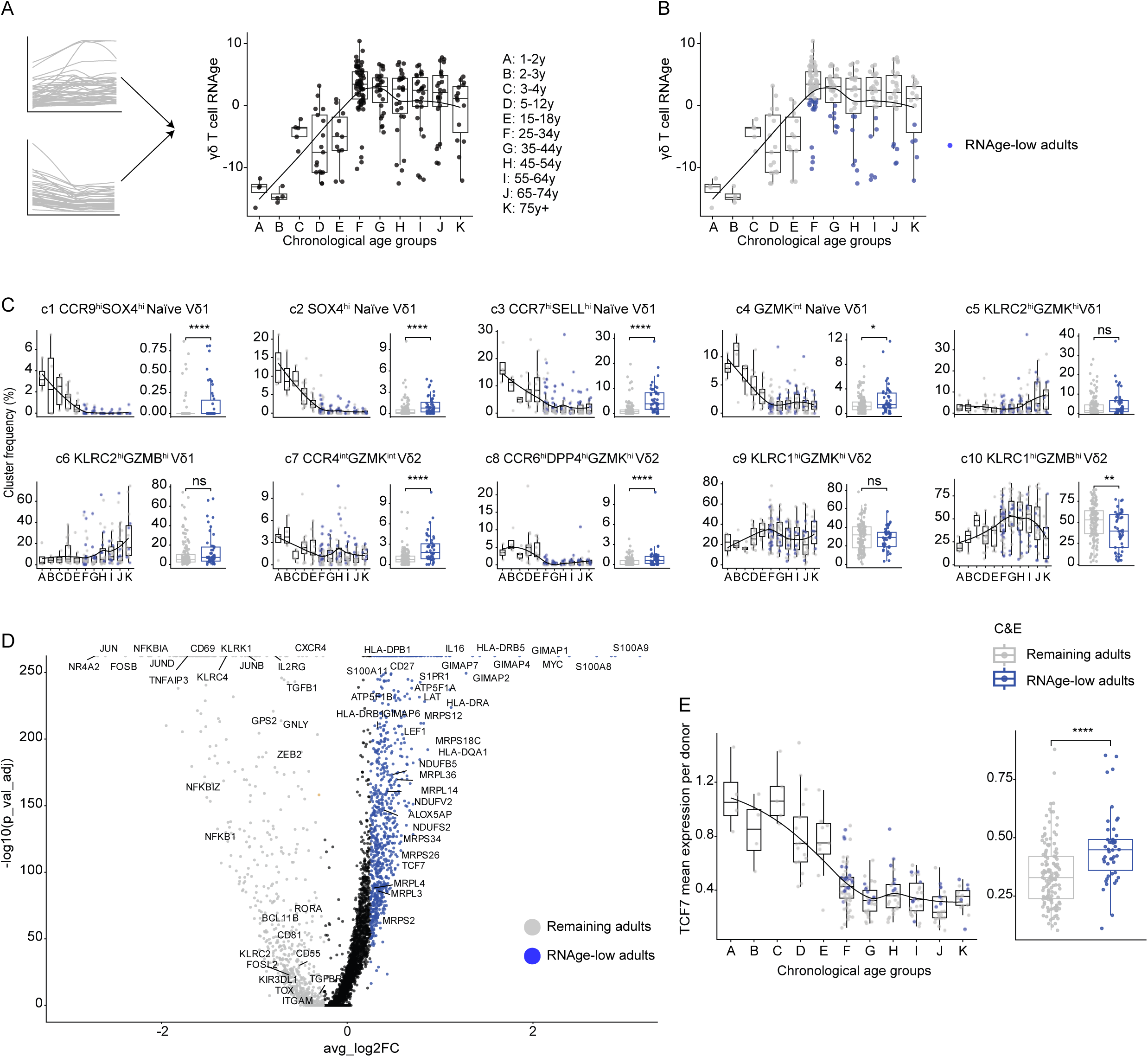
γδ T cell RNAge assesses age-related transcriptional programs and captures inter-individual heterogeneity. (**A**) γδ T cell RNA-based aging score (RNAge) was established using a gene set (**Supplemental Table 2**) associated with age-related transcriptional changes and assessed by principal component analysis at the donor level. (**B**) Box plot highlights adults with low RNAge scores, defined as individuals falling within the lowest quartile of each chronological age group. (**C**) Left, box plots display the frequencies of each γδ T-cell cluster across 11 age groups, with RNAge-low adults shown in blue. Right, comparison of cluster frequencies between RNAge-low adults and remaining adults. (**D**) Volcano plot shows differential gene expression between RNAge-low adults (blue) and remaining adults (grey). (**E**) Left: mean TCF7 expression per donor across age groups. Right: comparison of TCF7 expression between RNAge-low adults and remaining adults. Statistical comparison was assessed by unpaired two-sided Wilcoxon rank-sum test. Significance is denoted as ns (not significant), *p < 0.05, **p < 0.01, ***p < 0.001, ****p < 0.0001.

In addition, substantial inter-individual heterogeneity in γδ T cell RNAge was observed among adult donors (**Fig. 5A**). Adults whose RNAge score fell within the lowest quartile among individuals of each chronological age group were operationally defined as RNAge-low adults (**Fig. 5B**). Compared with remaining adults, RNAge-low adults displayed higher frequencies of naïve γδ T cell clusters c1-c4 as well as CCR4^int^ and CCR6^hi^ clusters c7 and c8, and decreased *GZMB*^hi^ Vδ2 cluster c10 (**Fig. 5C**).

Next, differential gene expression analysis between RNAge-low adults and remaining adults (chronological age groups F–K) was performed. This revealed increased expression of genes associated with T cell differentiation and stemness (e.g., *TCF7*, *LAT*, *CD27*, and *LEF1*) and mitochondrial activity (e.g., *NDUFB5*, *MRPL36*, and *ATP5F1A*) alongside reduced expression of effector and cytotoxicity-associated genes (e.g., *KLRC2*, *KLRC4*, *KLRK1*, *CD69*, and *GNLY*) in RNAge-low individuals (**Fig. 5D-E**). Notably, the transcription factor *TCF7*, which is critical for maintaining T cell stemness and regenerative potential, has been reported as a marker associated with immune resilience^47,48^. In addition, upregulation of multiple genes of the S100A and GIMAP families was observed in RNAge-low individuals.

Together, these results indicate that γδ T cell RNAge consolidates transcriptional changes of age and uncovers heterogeneous adult γδ T cell states that are not explained by chronological age alone.

## Discussion

Immune aging at the cellular level is often implicitly viewed as a gradual and continuous process across the lifespan, typically using linear models and largely inferred from studies of abundant immune cell populations and adult-derived PBMC cohorts^10,12,49^. γδ T cells, although present at low frequency in peripheral blood, represent a conserved third way of protection in human^50^. Here, by constructing a comprehensive single-cell transcriptome atlas of human γδ T cells across the entire lifespan, our study reveals that γδ T cell aging is not a linear lifelong process, but instead is characterized by pronounced remodeling in infancy and childhood, followed by long-term stability in adulthood.

At the population level, naïve Vδ1 T cells undergo a marked contraction while effector Vδ1 T cells expand with increasing age. In parallel, Vδ2 T cells peaked in childhood and declined in late adulthood, aligning with previous flow cytometry studies^26,30–32^. Moreover, γδ T cell gene expression profiles undergo a transition from high developmental gene expression signatures to expression of genes signatures linked to activation, cytotoxicity, inflammation, and anti-apoptosis. This shift is also accompanied by reduced mitochondrial-related gene expression, a hallmark of immunosenescence and inflammaging^51–54^. Importantly, both compositional and transcriptional changes are mainly established until late childhood, and show minimal changes throughout adulthood.

These observations are consistent with accumulating evidence indicating that aging involves sudden, accelerated, and non-linear transitions that often happen around midlife (40s) ^2,3^. However, our data indicate that major inflection of γδ T cell aging appears earlier, around young adulthood (20s). A possible explanation is their unique developmental origin and maturation program. As one of the earliest T cells developed in the fetal thymus, γδ T cells acquire effector programs, including the GZMA and GZMK production, and undergo expansion shortly after birth, accompanied by the acquisition of cytotoxic molecules such as GZMB, Perforin, and NKG2D ^28,29,55^. This also explains the limited age-related compositional and transcriptional variation of human γδ T cells from a previous single-cell transcriptomic study restricted to an adult cohort^16^.

In this context, we developed γδ T cell RNA age score (RNAge), which enables the assessment of overall γδ T cell transcriptional aging dynamics. Moreover, our analysis reveals substantial inter-individual heterogeneity among adults, which was not fully explained by chronological age. Notably, RNAge-low adults enriched for less differentiated features, characterized by increased naïve subsets and fetal-preferential subsets, reduced GZMB⁺ subsets, as well as enrichment of gene signatures for infant γδ T cells. Multiple factors such as genetic background, environments, diets, lifestyle, and infections (e.g., cytomegalovirus) are likely to contribute to inter-individual immune variations ^56^. However, the heterogeneous and limited availability for γδ T cell aging study precludes direct inference of drivers for γδ T cell RNAge variation. In addition, although cord blood datasets from our previous research are available ^41^, they were not included due to known immunological differences between cord blood and neonatal peripheral blood ^57–59^, limiting resolution at the earliest postnatal period. Nevertheless, γδ T cell RNAge provides a potential framework for assessing γδ T cell biological aging and identifying inter-individual heterogeneity.

Additionally, our analysis also provides insights into γδ T cell differentiation and effector maturation across age. We uncovered a *CCR9*^hi^*SOX4*^hi^ effector precursor subset that bifurcated into tissue-migratory and granzyme-producing effector lineages. The age-related attrition of this subset suggests progressive narrowing of the developmental reservoir for Vδ1 T cells in early childhood. In addition, within the γδ T cell effector pool, we described a clear transition from GZMK^+^ to GZMB^+^ perforin^+^ cells, mirroring maturation trajectories described in CD8⁺ T cells, MAIT and iNKT cells ^16,60^. Our study extends this concept to γδ T cells and, importantly, provides protein-level validation. In adults, GZMK⁺ cells comprise nearly half of the γδ T cell pool. Given the growing evidence linking GZMK⁺ cells to inflammaging, chronic inflammatory diseases and cancer^16,61–65^, the large GZMK⁺ pool in adults may serve as a reservoir for replenishing cytotoxic effectors and as a potential driver of chronic inflammation. Redirecting the GZMK→GZMB axis may therefore offer a therapeutic strategy to repurpose these pro-inflammatory intermediates toward protective cytotoxicity during aging.

In summary, we present a comprehensive single-cell transcriptional atlas of human γδ T cells from infancy to the elderly, establishing γδ T cells as a paradigm for non-linear immune aging at the cellular level. Furthermore, the *γδ T cells aging atlas* is fully accessible, offering a resource for future exploration and hypothesis generation.

## MATERIALS AND METHODS

### Study populations and Ethical approval

#### Healthy donors used for scRNA-seq

Details of included donors are provided in **Supplemental Table 1**. Recruitment and sample collection were conducted in accordance with the Declaration of Helsinki and approved by the institutional ethics board of Hannover Medical School (no. 10198_BO_S_2022, 8615_BO_S_2019). Samples from healthy toddlers and teenagers were collected at the Department of Pediatric Cardiology and Intensive Care Medicine and the Institute of Immunology, Hannover Medical School (Hannover, Germany). Healthy elderly donors were recruited as part of the RESIST Senior Individuals (SI) cohort ^66^.

#### Healthy CBMC and PBMC for flow cytometric analysis and in vitro stimulation

Recruitment and sample collection were performed under approval from the institutional ethics board of Hannover Medical School (no. 1303-2012, 3639-2017). Cord blood samples from healthy newborns were obtained from uncomplicated full-term pregnancies at Hannover Medical School. Peripheral blood from healthy adults was obtained from blood donors at the Department of Transfusion Medicine, Hannover Medical School.

### Isolation of lymphocytes

Peripheral blood mononuclear cells (PBMCs) were isolated from EDTA blood following density medium centrifugation. Isolated cells were gently frozen in a freezing medium containing 90% heat-inactivated fetal calf serum (FCS) and 10% dimethyl sulfoxide (DMSO), and stored in aliquots at −80 °C until further processing.

### scRNA-seq library construction from flow-sorted γδ T cells of healthy donors

Thawed PBMCs were stained with anti-CD3 (UCHT1, BioLegend), anti-γδ TCR BV510 (REA591, Miltenyi Biotec), anti-αβ TCR FITC (WT31, BD Biosciences), anti-CD56 APC (NCAM16.2, BD Biosciences), and anti-CD16 PerCP-Vio770 (REA423/3G8, Miltenyi Biotec). Dead cells were excluded by DAPI or Zombie Violet (BioLegend) staining.

Single-cell suspensions were subjected to library preparation using Chromium Next GEM Single Cell V(D)J Reagent Kits (10x Genomics). Depending on the donor cohort, two kit versions were employed: the v2 kit for toddlers, children, teenagers, and young adults, and the v1.1 kit for elderly donors. The experimental strategies differed across donor groups (**Supplemental Table 1**). For Healthy elderly (HealthyElderly1–12), each donor sample was labeled with a TotalSeq™ anti-human hashtag antibody (LNH-94, 2M2, BioLegend) prior to library preparation. Demultiplexing was performed with Souporcell ^67^, with hashtag information additionally incorporated for elderly samples. Donor sex was determined from the expression of Y chromosome–encoded genes (*RPS4Y1*, *DDX3Y*, *PRKY*, *UTY*, *ZFY*, *USP9Y*, *KDM5D*, *TMSB4Y*, *EIF1AY*).

Paired gene expression and surface protein libraries were sequenced on a NovaSeq X platform, targeting 25,000 and 5,000 read pairs per cell, respectively. Sequencing reads were aligned to the GRCh38 reference genome, and gene expression matrices were generated using Cell Ranger 7.0 (10x Genomics). Downstream analyses were conducted in R 4.4.2 using Seurat 5.2.0 ^68^.

### Computational extraction of γδ T cell subsets

For datasets sequenced together with αβ T cells, γδ T cells were identified based on the expression of *TRDV1*, *TRDV2*, *TRDV3*, and *TRDC*. For datasets containing both αβ T and NK cells, γδ T cells were identified using the same TCRδ gene criteria and by excluding cells expressing *NCAM1* (CD56). *FCGR3A*⁺ (CD16⁺) NK cell clusters were further removed to obtain a pure γδ T cell dataset.

### Integration of newly generated and published γδ T cell datasets

We re-analyzed previously published PBMC scRNA-seq datasets ^16,17^ and extracted γδ T cells according to the original annotations and computational extraction as described above according to the expression of *TRDV1*, *TRDV2*, *TRDV3*, and *TRDC* ^39^. Only donors with more than 100 γδ T cells were included in the analysis. Additional infant and aging single-cell cohorts were also examined (data not shown), but γδ T cell numbers in these datasets were insufficient for meaningful validation ^69–72^.

These re-processed γδ T cell datasets, together with the newly generated γδ T cell datasets from this study and γδ T cell datasets from our previous studies, were merged with the newly generated dataset using the Seurat merge function. Batch effects were corrected with the Harmony^40^ package (RunHarmony) using “batch” and “resource” metadata, followed by dimensionality reduction (RunUMAP) and clustering (FindClusters).

### Pseudotime analysis of Vδ1 T cells

Vδ1 T cells (clusters c1–c6) were isolated from the healthy γδ T cell atlas in Seurat and re-embedded with UMAP (20 Harmony-corrected PCs).

For Monocle 3 ^43^, the Seurat object was converted to a cell data set (as.cell_data_set) and analyzed following the standard Monocle 3 workflow. Cluster c1, corresponding to CCR9⁺SOX4⁺ Vδ1 cells, was specified as the root state. Principal graph–based pseudotime values were extracted, and for each trajectory, cells were stratified by donor age (groups A–L). Age-specific pseudotime distributions were visualized by Gaussian kernel density estimation (200 evaluation points, Silverman’s rule bandwidth), with densities normalized to unit area to allow direct comparison across age groups.

For Slingshot ^44^, UMAP coordinates and Seurat cluster labels were used as input, with cluster c1 as the start point. Two biologically consistent trajectories were retained.

### Differential gene expression (DEG) analysis between age groups

For age-associated comparisons, cells from age groups A–C were pooled as Young, and I–K as Old. Differentially expressed genes (DEGs) were identified using Seurat’s FindMarkers function (min.pct = 0.1, logfc.threshold = 0.1). Genes were assigned to “Young” or “Old” clusters based on the sign of avg_log2FC, and further filtered to retain those with |avg_log2FC| > 0.5, an absolute difference in detection rate between groups (|pct.1 − pct.2| > 0.1), and exclusion of ribosomal (RP*) and mitochondrial (MT*) genes. Module scores for selected gene sets were computed using Seurat’s AddModuleScore function.

### Gene Ontology (GO) enrichment analysis

Functional enrichment of DEGs was performed using the compareCluster function in the clusterProfiler^73^ R package (enrichGO, ontology = Biological Process, OrgDb = *org.Hs.eg.db*, keyType = “SYMBOL”, pAdjustMethod = “BH”, qvalueCutoff = 0.05). Gene ratios were calculated as the percentage of DEGs annotated to each term relative to the total background genes. Selected GO terms of interest were visualized using ggplot2.

### Flow cytometric analysis

For GZMK and GZMB staining, CBMCs and PBMCs were thawed in a 37 °C water bath, counted, and 0.5 - × 10⁶ cells were used for antibody staining. Cells were stained for 30 min at room temperature with the following surface antibodies: anti-CD3 AF532 (17A2, eBioscience), anti-γδ TCR PE (11F2, Miltenyi Biotec), anti-CD19 APC-Fire810 (HIB19, BioLegend), anti-Vδ2 APC-Vio770 (15D, Miltenyi Biotec), anti-Vδ1 VioGreen (TS8.2, Miltenyi Biotec), anti-CD16 BUV496 (3G8, BD Biosciences), anti-CD69 BUV737 (FN50, BD Biosciences), anti-CD127 BV785 (A019D5, BioLegend), anti-CD28 PE-Cy5 (CD28.2, BioLegend), and anti-CD27 AF700 (O323, BioLegend). Dead cells were excluded using Zombie NIR (BioLegend). Following surface staining, cells were fixed and permeabilized using the Foxp3/Transcription Factor Staining Buffer Set (eBioscience) according to the manufacturer’s protocol, and stained intracellularly with anti-Perforin BV421 (dG9, BioLegend), anti-GZMK PerCP-eFluor710 (GM26E7, Invitrogen), and anti-GZMB AF647 (GB11, BioLegend).

### FACS sorting of CD28^+^ Vδ2 T cells and HMBPP stimulation

PBMCs were thawed, counted, and ∼2 × 10⁷ cells per donor were stained for 15 min at room temperature with anti-CD3 AF488 (UCHT1, BioLegend), anti-γδ TCR PE (11F2, Miltenyi Biotec), anti-CD28 PE-Cy5 (CD28.2, BioLegend), and anti-Vδ2 APC-Vio770 (15D, Miltenyi Biotec). Dead cells were excluded with Zombie Violet (BioLegend). CD28⁺ Vδ2 T cells (live CD3⁺ γδTCR⁺ Vδ2⁺) and non-T cells (live CD3⁻) were sorted on a FACSAria III (BD Biosciences).

For baseline phenotyping (day 0), ∼1 × 10⁴ sorted CD28⁺ Vδ2 T cells and 5 × 10^5^ PBMCs (staining control) were stained with the following antibodies: anti-CD16 BUV496 (3G8, BD Biosciences), anti-Perforin BV421 (dG9, BioLegend), anti-Vδ1 VioGreen (TS8.2, Miltenyi Biotec), anti-CD3 AF488 (UCHT1, BioLegend), anti-GZMK PerCP-eFluor710 (clone GM26E7, Invitrogen), anti-GZMB AF647 (GB11, BioLegend), anti-CD27 AF700 (O323, BioLegend), and anti-Vδ2 APC-Vio770 (15D, Miltenyi Biotec).

For stimulation assays, ∼3 × 10⁴ sorted CD28⁺ Vδ2 T cells were co-cultured with non-T cells at a 1:10 ratio in the presence of 100 U/mL human IL-2 (Sigma-Aldrich) and 1 μM HMBPP (Sigma-Aldrich). As staining controls, 0.5 × 10⁶ PBMCs from each donor were cultured with 100 U/mL IL-2 alone. After 3 days, cells were stained using the same protocol and antibody panel as for day 0, followed by acquisition on the Cytek Aurora as described above.

### γδ T cell RNAge establishment, RNAge-low adult identification, and differential gene expression analysis

γδ T cell RNA-based aging score (γδ T cell RNAge) was generated using a gene set showing age-related changes identified in Figure 4 (**Supplemental Table 2**). For each cell, normalized expression values of all genes were extracted from the Seurat object. Genes increasing with age were multiplied by −1. Values were subsequently aggregated at the donor level by calculating the mean expression across all γδ T cells per donor. Principal component analysis (PCA) was then performed on the donor-level expression matrix (scaled), and the first principal component (PC1) was used as the γδ T cell RNAge score, capturing the dominant axis of age-associated transcriptional variation.

Next, to identify adults with relatively low γδ T cell RNA aging profiles, analysis was restricted to adult donors (age groups G–L). A LOESS regression model was fitted to describe the expected relationship between chronological age groups and γδ T cell RNAge score. For each donor, the residual was calculated as the difference between the observed RNAge score and the LOESS-fitted value. Adult donors with residuals below the 25^th^ percentile were classified as RNAge-low adults, whereas donors within the interquartile range were considered typical.

Differential gene expression analysis was performed using Seurat’s FindMarkers function (min.pct = 0.1, logfc.threshold = 0.1). DEGs were further filtered to retain those with |avg_log2FC| > 0.5, an absolute difference in detection frequency between groups (|pct.1 − pct.2| > 0.1), and exclusion of ribosomal (RP*) and mitochondrial (MT*) genes.

### Statistical analysis

Comparisons of γδ T cell cluster frequencies and flow cytometry measurements between independent groups were performed using the unpaired two-sided Wilcoxon rank-sum test. Paired comparisons of in vitro stimulation experiments were assessed by the paired two-sided Wilcoxon signed-rank test. For scRNA-seq analyses, differential gene expression was identified using Seurat’s FindMarkers function as described above, with significance determined by adjusted p < 0.05. Unless otherwise indicated, significance thresholds were set as ns (not significant), p < 0.05, *p < 0.01, **p < 0.001, and ***p < 0.0001. No correction for multiple testing was applied to flow cytometry–based comparisons due to the exploratory nature of this study. All analyses were conducted in R 4.4.2.

## Acknowledgments

We thank the RESIST Senior Individuals cohort investigators for providing resources. We thank Dr. Matthias Ballmaier and the sorting facility at Hannover Medical School for their kind support in performing FACS. We thank Prof. Britta Eiz-Vesper and the Institute of Transfusion Medicine and Transplant Engineering of Hannover Medical School to support the project with resources.

## Author contributions statement

Study concept and design: T.Y., S.R.; Resources: X.L-L, R.F., C.v.K., I.P.; experiments: T.Y., C.K., A.J., V. A., Y. A., Z.W., M.D.; data analysis: T.Y., A.H., Z.S., L.T.; data interpretation: T.Y., S.R.; data visualization: T.Y.; statistics: T.Y.; writing of the manuscript: T.Y., S.R. The manuscript was read and approved by all authors.

## Data and materials availability

Raw FASTQ files are deposited in the NCBI Gene Expression Omnibus (GSE306534 and GSE306377). Processed scRNA-seq expression matrices, including metadata and normalized counts for generating Seurat object RDS files of the healthy γδ T cell atlas will be available at https://www.synapse.org.

This study did not generate new code and default parameters were used for all arguments unless otherwise mentioned. The scripts used to generate Seurat objects for the healthy γδ T cell aging atlas will be available at https://www.synapse.org.

## Data and code Availability

Raw FASTQ files are deposited in the NCBI Gene Expression Omnibus (GEO) under accessions GSE306534 and GSE306377.

## Supplementary Materials

Fig. S1 to S3 for multiple supplementary figures.

Table S1 to S2 for multiple supplementary tables.

