## supplemental information for "Transcriptomic profiling of human γδ T cells reveals non-linear immune aging characterized by childhood transitions and relative stability in adulthood"

### Supplementary Table 1. Metadata of healthy donors included in single-cell transcriptomic analysis.

| Basic | | | | | | | | |
|---|---|---|---|---|---|---|---|---|
| SampleID | OriginalID | Sex | Age groups | CMVsero Status | 10x Chromium Next GEM Single Cell 5' Kit | 10x Genomics Reaction Setup | Demultiplexing by | donorID in Seurat object |
| HealthyToddler1 | HC 007 | m | D_3-4y | Unknown | v2 | One 10x Reaction | Souporcell | 3y_0, 3y_1, 3y_2, 3y_3, 3y_4 |
| HealthyToddler2 | HC 008 | m | D_3-4y | Unknown | v2 | | Souporcell | |
| HealthyToddler3 | HC 023 | m | D_3-4y | Unknown | v2 | | Souporcell | |
| HealthyToddler4 | HC 005 | m | D_3-4y | Unknown | v2 | | Souporcell | |
| HealthyToddler5 | HC 015 | unkown | D_3-4y | Unknown | v2 | | Souporcell | |
| HealthyChildren1 | HC045 | unkown | E_5-12y | Sero positive | v2 | One 10x Reaction | Souporcell | ChildrenCMV_0, ChildrenCMV_1,ChildrenCMV_2 |
| HealthyChildren2 | HC054 | unkown | E_5-12y | Sero positive | v2 | | Souporcell | |
| HealthyChildren3 | HC051 | unkown | E_5-12y | Sero positive | v2 | | Souporcell | |
| HealthyTeenager1 | HC 018 | unkown | F_14-18y | Unknown | v2 | One 10x Reaction | Souporcell | 15y_0, 15y_1, 15y_2, 15y_3, 15y_4, 15y_5 |
| HealthyTeenager2 | HC 009 | unkown | F_14-18y | Sero negative | v2 | | Souporcell | |
| HealthyTeenager3 | HC 038 | unkown | F_14-18y | Sero negative | v2 | | Souporcell | |
| HealthyTeenager4 | HC 047 | unkown | F_14-18y | Unknown | v2 | | Souporcell | |
| HealthyTeenager5 | HC 002 | unkown | F_14-18y | Unknown | v2 | | Souporcell | |
| HealthyTeenager6 | HC 039 | F | F_14-18y | Unknown | v2 | | Souporcell | |
| HealthyYoungAdult1 | TMHC14 | F | G_25-34y | Sero positive | v2 | One 10x Reaction | Souporcell | A_CMV_0, A_CMV_1 |
| HealthyYoungAdult2 | TMHC17 | F | G_25-34y | Sero positive | v2 | | Souporcell | |
| HealthyElderly1 | DRESD000019 | F | K_65-74y | Sero positive | v1 | One 10x Reaction | Hashtag and Souporcell | CMV_1_1_019 |
| HealthyElderly2 | DRESD000094 | M | L_75y+ | Sero positive | v1 | | Hashtag and Souporcell | CMV_1_2_094 |
| HealthyElderly3 | DRESD000129 | M | K_65-74y | Sero positive | v1 | One 10x Reaction | Hashtag and Souporcell | CMV_2_1_129 |
| HealthyElderly4 | DRESD000079 | M | K_65-74y | Sero positive | v1 | | Hashtag and Souporcell | CMV_2_3_079 |
| HealthyElderly5 | DRESD000105 | M | L_75y+ | Sero positive | v1 | | Hashtag and Souporcell | CMV_2_2_105 |
| HealthyElderly6 | DRESD000107 | F | K_65-74y | Sero positive | v1 | | Hashtag and Souporcell | CMV_2_0_107 |
| HealthyElderly7 | DRESD000141 | F | K_65-74y | Sero positive | v1 | One 10x Reaction | Hashtag and Souporcell | CMV_3_1_141 |
| HealthyElderly8 | DRESD000070 | M | K_65-74y | Sero positive | v1 | | Hashtag and Souporcell | CMV_3_2_070 |
| HealthyElderly9 | DRESD000370 | F | J_55-64y | Sero positive | v1 | | Hashtag and Souporcell | CMV_3_2_370 |
| HealthyElderly10 | DRESD001344 | F | K_65-74y | Sero positive | v1 | One 10x Reaction | Hashtag and Souporcell | CMV_4_1_344 |
| HealthyElderly11 | DRESD000339 | M | K_65-74y | Sero positive | v1 | | Hashtag and Souporcell | CMV_4_2_339 |
| HealthyElderly12 | DRESD000331 | F | L_75y+ | Sero positive | v1 | | Hashtag and Souporcell | CMV_4_0_331 |

**Supplemental Table 2: genes used in Figure 5**

```
age_down_genes <- c(
  "LEF1","CD27","TCF7","LTB","ID3","CCR7","SOX4","MYC","CCR9","LAT",
  "NDUFV2","NDUFS2","NDUFB1","NDUFB4","NDUFB5","UQCRC1","UQCRC2","UQCC2",
  "COX7A2L","ATP5F1A","ATP5F1B","ATP5F1C","ATP5MF","ATP5ME","STOML2",
  "OXA1L","NDUFAF4","MRPS2","MRPS5","MRPS6","MRPS12","MRPS18B","MRPS18C",
  "MRPS26","MRPS34","MRPL3","MRPL4","MRPL14","MRPL17","MRPL28","MRPL34",
  "MRPL36","MRPL51","MRPL52"
)

age_up_genes <- c(
  "HLA-DRB5", "HLA-DPB1", "HLA-DRA", "HLA-DQA1", "HLA-DRB1",
  "CD160", "ZEB2", "CD81", "CD69",
  "ITGA4", "ITGAM", "ITGB1",
  "LGALS1", "CX3CR1", "CXCR4", "S1PR5",
  "CDC42", "PREX1", "TMSB10",
  "NFKBIZ", "ZBP1", "ALOX5AP", "GPS2", "CTSC",
  "TNFRSF1B", "FGR", "CARD16", "CASP1", "TNFAIP3", "TGFB1", "RORA",
  "CST7", "CD53", "TYROBP", "NFKB1",
  "KIR3DL1", "KLRC1", "KLRB1", "KLRC4",
  "GZMM", "FCGR3A", "KLRK1", "NKG7",
  "PRF1", "GZMB", "KLRD1", "GZMH", "GNLY", "KLRG1",
  "CCL5",
  "JUN", "JUNB", "JUND",
  "FOS", "FOSB", "FOSL2"
)
```

Figure S1

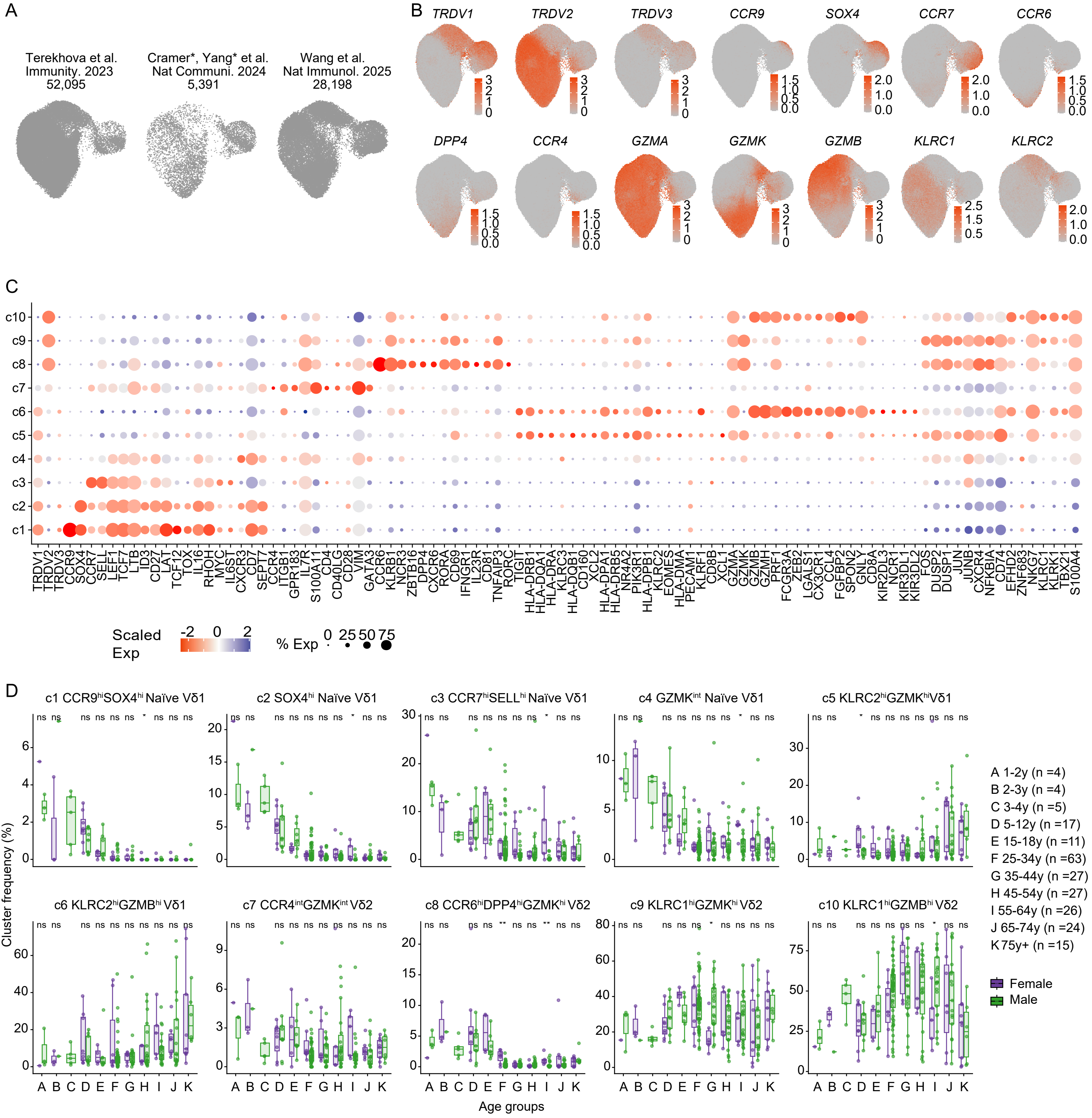


**Figure S1: Single-cell transcriptome analysis of human γδ T cells from newborns to elderly, related to Figure 1.**
(**A**) UMAP plots showing public datasets used in this study. (**B**) UMAP plots showing expression of selected marker genes at the single-cell level. (**C**) Dot plot showing scaled expression (Scaled Exp) and proportion (% Exp) of cells expressing selected highly expressed genes across clusters c1–c10. (**D**) Cluster frequencies (c1–c10) across 12 age groups (A–K), stratified by sex. Each dot represents one donor; boxes indicate median and interquartile range. Male and female donors are indicated by color (purple = female, gray = male). Statistical comparisons were assessed by unpaired two-sided Wilcoxon rank-sum test. Significance is denoted as ns (not significant), *p < 0.05, **p < 0.01, ***p < 0.001, ****p < 0.0001.

Figure S2

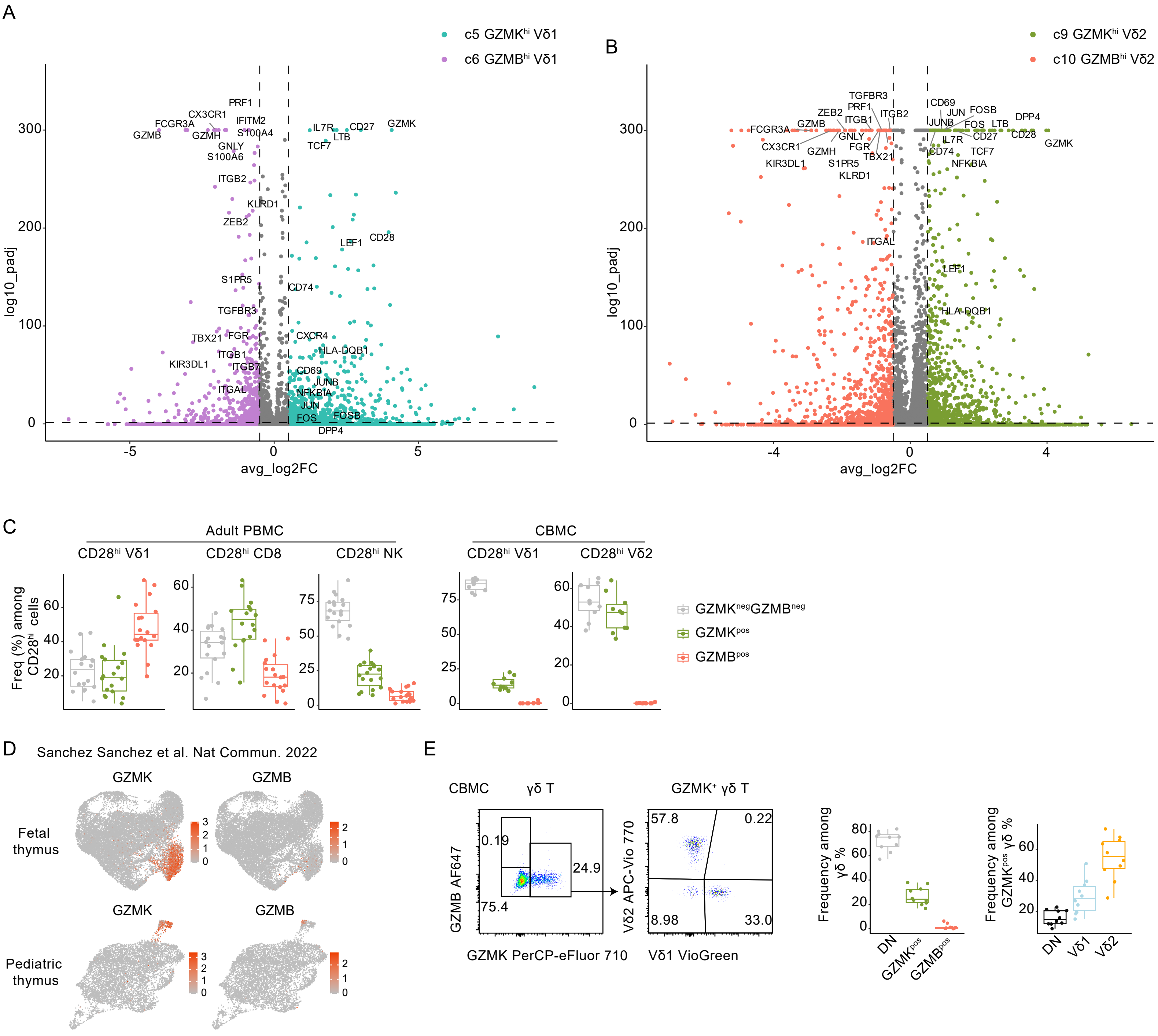


**Figure S2: Characterization of GZMK$^+$ and GZMB$^+$ γδ T cell subsets, related to Figure 3.**
(**A-B**) Volcano plots showing differentially expressed genes (DEGs) between GZMK$^+$ and GZMB$^+$ Vδ1 (**A**) and Vδ2 (**B**) T cells. Selected genes are labeled. (**C**) Frequencies of GZMK$^+$GZMB$^+$, GZMK$^-$GZMB$^-$, and GZMK$^+$GZMB$^+$ subsets among CD28$^+$ γδ T cells, CD8$^+$ T cells, and NK cells from adult PBMCs (n = 18), and among CD28$^+$ Vδ1$^+$ and Vδ2$^+$ γδ T cells from CBMCs (n = 10). Each dot represents one donor. Boxes indicate median and interquartile range. (**D**) UMAP plots showing GZMK and GZMB expression in γδ T cells from human fetal and pediatric thymus (re-analyzed from Sanchez Sanchez et al., Nat Commun 2022). (**E**) Flow cytometry analysis of CBMC-derived γδ T cells (n = 10). Left: Representative gating of GZMK and GZMB expression in total γδ T cells and representative Vδ1 and Vδ2 expression in GZMK$^+$ γδ T cells. Right: Frequencies of DN, GZMK$^+$, and GZMB$^+$ subsets among total γδ T cells, and Vδ1, Vδ2, and DN subsets among GZMK$^+$ γδ T cells. Each dot represents one donor; boxes indicate median and interquartile range.

Figure S3-1

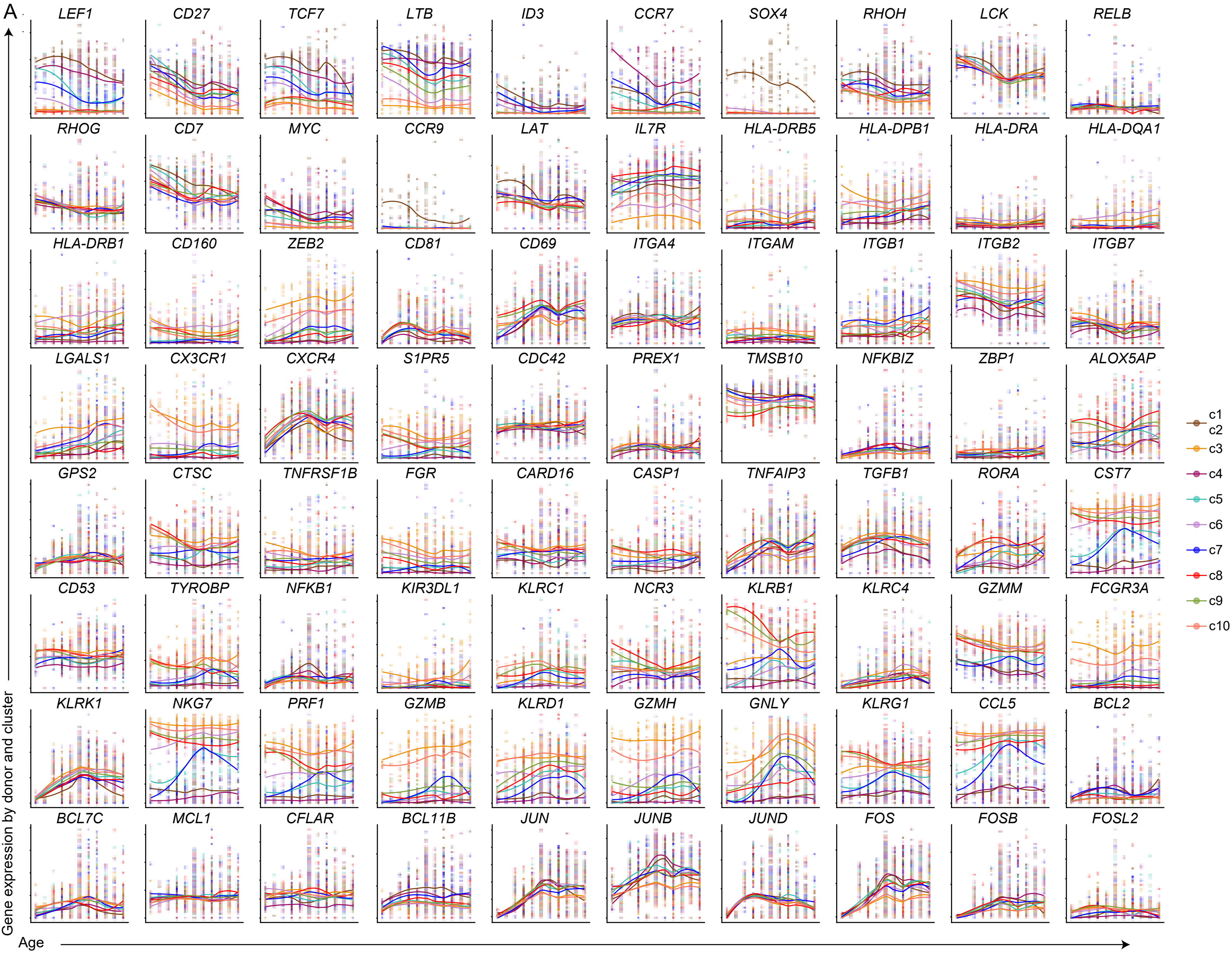


**Figure S3: Age-related expression dynamics of selected genes across γδ T cell clusters, related to Figure 4.**
(**A**) Gene expression trends of selected developmental, activation, migration, cytotoxicity, and transcription factor genes plotted across age and cluster identity. The scores were computed with defined gene sets (γδ T cell development: *LEF1, CD27, TCF7, LTB, ID3, CCR7, SOX4, RHOH, LCK, RELB, RHOG, CD7, MYC, CCR9, LAT, IL7R*; Activation: *HLA-DRB5, HLA-DPB1, HLA-DRA, HLA-DQA1, HLA-DRB1, CD160, ZEB2, CD81, CD69*; Migration: *ITGA4, ITGAM, ITGB1, ITGB2, ITGB7, LGALS1, CX3CR1, CXCR4, S1PR5, CDC42, PREX1, TMSB10;* Inflammatory response: *NFKBIZ, ZBP1, ALOX5AP, GPS2, CTSC, TNFRSF1B, FGR, CARD16, CASP1, TNFAIP3, TGFB1, RORA, CST7, CD53, TYROBP, NFKB1*; Cell cytotoxicity: *KIR3DL1, KLRC1, NCR3, KLRB1, KLRC4, GZMM, FCGR3A, KLRK1, NKG7, PRF1, GZMB, KLRD1, GZMH, GNLY, KLRG1, CCL5*; Anti-apoptosis: *BCL2, BCL7C, MCL1, CFLAR, BCL11B; AP-1* Transcrition factors: *JUN, JUNB, JUND, FOS, FOSB, FOSL2*). (**B**) Expression dynamics of genes related to cytokines and cytokine receptors, chemokines, TCR signaling components, toll-like receptors, and proliferation markers, plotted across age. (**C**) Expression patterns of genes associated with canonical hallmarks of immune aging, including T cell exhaustion, SASP (senescence-associated secretory phenotype), DNA repair and genome maintenance, autophagy, and epigenetic regulation. Gene expression values are shown as donor-level averages per cluster. Curves represent LOESS-smoothed trends across age.

Figure S3-2

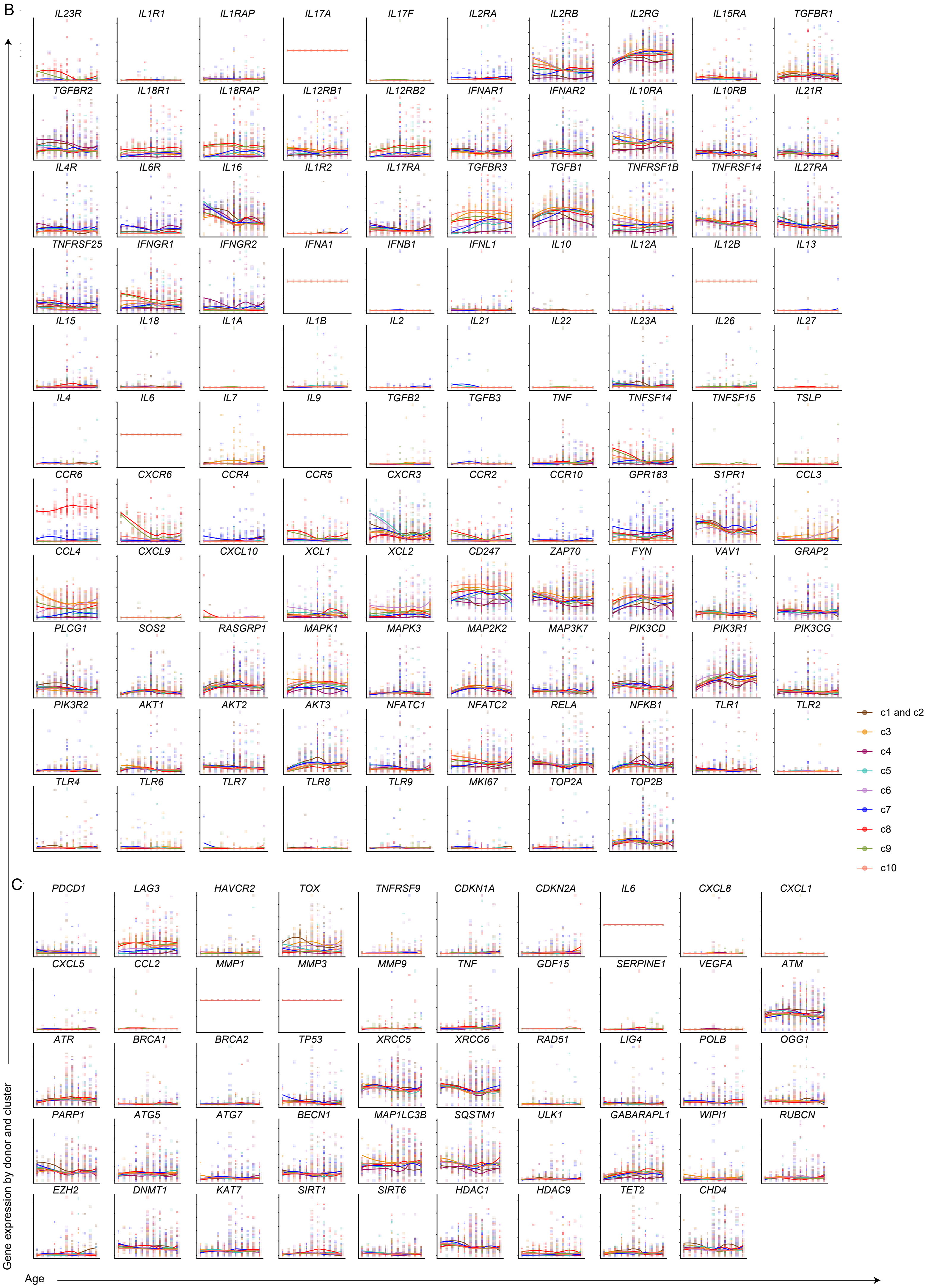

B
IL23R IL1R1 IL1RAP IL17A IL17F IL2RA IL2RB IL2RG IL15RA TGFBR1
TGFBR2 IL18R1 IL18RAP IL12RB1 IL12RB2 IFNAR1 IFNAR2 IL10RA IL10RB IL21R
IL4R IL6R IL16 IL1R2 IL17RA TGFBR3 TGFB1 TNFRSF1B TNFRSF14 IL27RA
TNFRSF25 IFNGR1 IFNGR2 IFNA1 IFNB1 IFNL1 IL10 IL12A IL12B IL13
IL15 IL18 IL1A IL1B IL2 IL21 IL22 IL23A IL26 IL27
IL4 IL6 IL7 IL9 TGFB2 TGFB3 TNF TNFSF14 TNFSF15 TSLP
CCR6 CXCR6 CCR4 CCR5 CXCR3 CCR2 CCR10 GPR183 S1PR1 CCL3
CCL4 CXCL9 CXCL10 XCL1 XCL2 CD247 ZAP70 FYN VAV1 GRAP2
PLCG1 SOS2 RASGRP1 MAPK1 MAPK3 MAP2K2 MAP3K7 PIK3CD PIK3R1 PIK3CG
PIK3R2 AKT1 AKT2 AKT3 NFATC1 NFATC2 RELA NFKB1 TLR1 TLR2
TLR4 TLR6 TLR7 TLR8 TLR9 MKI67 TOP2A TOP2B
c1 and c2
c3
c4
c5
c6
c7
c8
c9
c10
C
PDCD1 LAG3 HAVCR2 TOX TNFRSF9 CDKN1A CDKN2A IL6 CXCL8 CXCL1
CXCL5 CCL2 MMP1 MMP3 MMP9 TNF GDF15 SERPINE1 VEGFA ATM
ATR BRCA1 BRCA2 TP53 XRCC5 XRCC6 RAD51 LIG4 POLB OGG1
PARP1 ATG5 ATG7 BECN1 MAP1LC3B SQSTM1 ULK1 GABARAPL1 WIPI1 RUBCN
EZH2 DNMT1 KAT7 SIRT1 SIRT6 HDAC1 HDAC9 TET2 CHD4
Gene expression by donor and cluster
Age
